# Coupling of ATP and PMF in *Escherichia coli* is determined by growth conditions

**DOI:** 10.64898/2026.04.21.719851

**Authors:** Ekaterina Krasnopeeva, Bryan Wu, Stefan Kittler, Calin C. Guet

## Abstract

The ATP molecule is the universal energy currency across all living organisms. There are two fundamental pathways of ATP synthesis: substrate-level and oxidative phosphorylation. While substrate-level phosphorylation generates ATP directly, in oxidative phosphorylation, proton motive force (PMF) is required to power ATP synthesis via the F_1_F_o_ ATP synthase. Using *Escherichia coli*, we show that due to simultaneous use of both pathways, the strength of coupling between ATP and PMF strongly depends on growth conditions: coupling is weak when requirements for independent generation of ATP and PMF are met, and becomes essential when not. We determine the conditions, under which PMF-ATP coupling becomes essential and show that PMF is required for bacterial growth irrespective of its ATP synthesis function. We propose that the main role of F_1_F_o_ in *Escherichia coli*, contrary to the canonical view, is not to generate ATP but to provide an auxiliary pathway that allows both, ATP and PMF, to be produced.

## Introduction

Adenosine-5’-triphosphate (ATP) is a universal energy currency used by every known organism. Most energy-consuming processes in a living cell are coupled with the conversion of ATP to adenosine-5’-diphosphate (ADP) and inorganic phosphate (P_*i*_) to be energetically favourable.

There are two fundamental mechanisms by which organisms synthesise ATP: direct (substrate-level phosphorylation) and indirect (electron transport-linked phosphorylation) [1–3]. During substrate-level phosphorylation, the phosphate group is transferred from a high-energy substrate to ADP directly to form ATP. In the process of electron transport-linked phosphorylation, which includes both, oxidative phosphorylation and photophosphorylation, ATP synthesis is powered by an electrochemical gradient of protons, also known as proton motive force (PMF), via a chemiosmotic mechanism [1–5].

PMF and ATP are interconnected via the enzyme F_1_F_o_ ATP synthase/ATPase. In 1961 Peter Mitchell proposed that the energy stored in the form of PMF can be converted to ATP via F_1_F_o_ ATP synthase, where the flow of protons across the membrane down the gradient powers the ADP+P_*i*_ → ATP reaction [4, 5]. The reaction can be reversed: when working as ATPase, F_1_F_o_ hydrolyses ATP to ADP and P_*i*_, creating an outward proton flux and thereby generating PMF [6, 7].

In bacteria, specifically in *Escherichia coli*, two pathways of energy metabolism are usually considered: *fermentation* and *respiration* [1, 8]. We note here, that these terms are not synonymous to the fundamental mechanisms of substrate-level and oxidative phosphorylation we discussed earlier.

*Respiration* relies on the full oxidation of the substrate through a series of chemical reactions within glycolysis and tricarboxylic acid (TCA, or Krebs) cycle, yielding a certain amount of substrate-level produced ATP (the amount of ATP differs depending on a specific metabolic pathway for a given carbon source), and electron carriers: reduced nicotinamide adenine dinucleotide (NADH) and reduced flavin adenine dinucleotide (FADH_2_). NADH and FADH_2_ are then oxidised by components of the electron transport chain (ETC), transferring electrons to quinones and then to terminal oxidases, which in turn pump protons across the cytoplasmic membrane to generate PMF [1, 2, 8]. This process requires the presence of a terminal electron acceptor for NADH/FADH_2_ oxidation (oxygen in aerobic respiration; nitrate, fumarate, or other compounds in anaerobic respiration). In the absence of a terminal electron acceptor, NADH/FADH_2_ are accumulated in their reduced forms and stall the cycle, thereby inhibiting respiration [9, 10].

*Fermentation* does not require oxygen. It also produces substrate-level ATP and NADH. In the presence of oxygen, these NADHs can be converted to PMF via ETC, similar to respiration, allowing further oxidative phosphorylation. If no electron acceptor is present, the redox balance is maintained by using NADH to reduce organic intermediates instead, thereby regenerating NAD^+^ [8, 11]. Importantly, both fermentation and respiration can produce ATP by substrate-level and oxidative phosphorylation, if the requirements are met (fermentable carbon source and terminal electron acceptor are present).

Respiration is a more energy-efficient pathway, since it yields more ATP molecules per molecule of glucose (or other carbon source, see SI Table 1). However, fermentation is faster [12] and more efficient in proteome allocation, which is especially crucial for fast-growing cells [13, 14]. For over a century now it is known that tumour cells in human body prefer fermentative pathways over respiration even in the presence of oxygen – a phenomenon known as the Warburg effect [15–17]. Similar effects are observed in yeasts (Crabtree Effect) [18] and bacteria (overflow metabolism) [19, 20]. For bacteria, overflow metabolism describes the process where, while growing above a characteristic growth rate in the presence of oxygen, cells produce acetate, which indicates that both fermentation and respiration pathways are used at the same time [13, 20– 22].

In summary, in the presence of oxygen and a fermentable carbon source, *E. coli* can simultaneously metabolise through respiratory and fermentative pathways, each of which produces ATP both directly and indirectly via PMF. This suggests a highly intricate relationship between PMF and ATP, that are often assumed to be tightly coupled. Here, we study the strength of this coupling and its dependence on the growth conditions, as defined by different carbon sources and the availability of oxygen.

## Results

### In the presence of glucose uncouplers of oxidative phosphorylation do not cause ATP depletion in *Escherichia coli*

To determine the strength of ATP and PMF coupling, we first test how uncoupling of oxidative phosphorylation affects ATP levels in the cell cytoplasm. When uncoupled, ATP can no longer be produced via PMF-driven ATP synthase, but only via substrate-level phosphorylation. We choose to do it in the defined minimal medium M9, commonly used for *E. coli* growth under laboratory conditions, supplemented with 0.3% glucose (M9Glu). In this medium energy is produced solely from glucose metabolism. We grow cells in M9Glu to OD 0.3-0.5, split the culture in two, and wash each of the aliquots in either fresh M9Glu medium or in M9 medium containing no glucose, where no metabolism is possible. ATP measurements start immediately after the wash. We use three different uncouplers, most commonly used in the literature: two protonophores, carbonyl cyanide m-chlorophenyl hydrazone (CCCP) [23–25] and 2,4-dinitrophenol (DNP) [26, 27], as well as sodium azide (SA), which works as an ETC inhibitor [28, 29] and inhibitor of ATP hydrolysis by F_1_F_o_ ATPase [30–32].

Protonophores dissipate PMF by shuttling protons down their gradient across the lipid membrane, while ETC and ATPase inhibition fully stops PMF production. The initial level of ATP in the population is measured prior to the treatment with uncouplers (t=0). Then, 100 *µ*M CCCP, 5 mM SA, or 2.5 mM DNP is added to the culture and ATP level is measured every 5-10 minutes with luciferase assay for a total of 1 hour, Fig. 1A. To convert the total amount of ATP in a bacterial population to the ATP concentration in a single cell cytoplasm, the number of cells is estimated from the CFU count, Fig. 1B, and cell volume is estimated by phase contrast microscopy, Fig. SI 1. Interestingly, none of the treatments induces cell death, with the CFU number remaining roughly constant for the duration of the experiment, Fig. 1B.

**Fig. 1:**
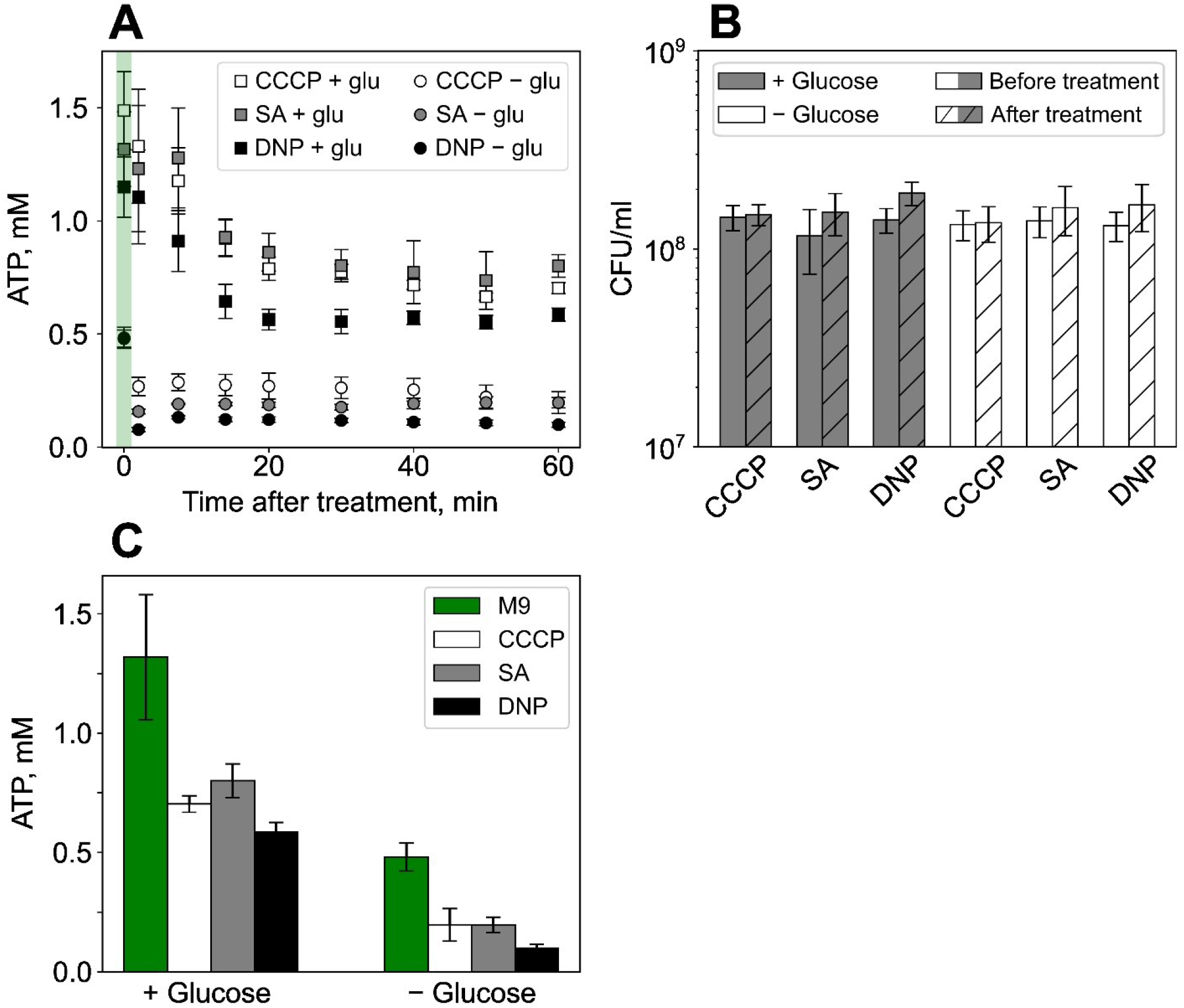
Uncouplers of oxidative phosphorylation reduce but do not deplete ATP in the cells if glucose is present in the medium. (**A**) Time series of ATP concentration in *E. coli* cytoplasm upon addition of PMF-ATP uncoupling agents – 100 *µ*M CCCP (white markers), 5 mM SA (grey markers), and 2.5 mM DNP (black markers) – in medium containing 0.3% glucose (squares) or no carbon source (circles). Uncouplers are added at time t=0 and ATP concentration is measured every 5-10 minutes for a total of 1 hour with luciferase assay. The first point at t=0 shows ATP concentration prior to treatment (highlighted by the green shading). Error bars show standard deviation of the mean for N=3 independent experiments for each treatment. (**B**) Absolute ATP concentration in cells’ cytoplasm before (t=0, green bars) or after (t=60 min) CCCP, SA or DNP addition, with and without glucose. (**C**) CFU counts before (t=0) and after (t=60 min) treatment. Within 60 minutes of the experiment CFU count stays constant or increases slightly indicating that cells neither grow (CFU increase corresponds to less than 1 division) nor die when uncouplers are present in the medium, irrespective of glucose.

We find that if glucose is present in the medium, ATP level decreases approximately by half from the initial 1.32 ± 0.26 mM to 0.70 ± 0.03 mM after 1 hour treatment with CCCP, 0.80 ± 0.03 mM with SA, and 0.59 ± 0.04 mM with DNP, Fig. 1C. Although this is a significant decrease in ATP concentration, its level remains relatively high. ATP level reduction happens gradually within approximately 20 minutes after an uncoupler introduction and remains constant for the rest of the experiment. In contrast, PMF is fully depleted immediately after 100 *µ*M CCCP addition, as we have shown previously [33]. With ATP consumption rate of ∼6 million molecules per second [34], ATP concentration in the cytoplasm ∼1 mM (Fig. 1C), and cell volume ∼2 femtoliters (Fig. SI 1), we can calculate the turnover rate for the whole ATP pool to be less than 1 second, so the gradual ATP concentration decrease cannot be explained by its normal consumption.

In contrast, in the medium with no glucose, cells start with significantly lower cytoplasmic ATP (0.48 ± 0.06 mM, Fig. 1C) and reach the new ATP level (0.20 ± 0.07 mM with CCCP, 0.20 ± 0.03 mM with SA, and 0.10 ± 0.02 mM with DNP) immediately after exposure to the uncouplers. Interestingly, the 10-15 minutes of starvation cells experience in glucose-deficient medium during the wash before the start of the ATP measurements cause their cytoplasmic ATP level to drop below the final ATP level after 60 minutes treatment with uncouplers in glucose-rich medium, Fig. 1C. However, ATP level drops even further, and immediately, when the uncouplers are added. We assume that the final drop is related to the residual metabolism on glucose left over after the wash.

High level of ATP after PMF-depleting treatments suggest that the large portion of ATP is being synthesised via non-PMF-dependent pathways, i.e., substrate-level phosphorylation. This result was somewhat surprising, since, theoretically, substrate-level phosphorylation accounts for less than 15% of total (theoretical maximum) ATP produced from glucose metabolism (see Fig. SI 2 and SI Table 1). Therefore, we decided to investigate further, how strongly *E. coli* depend on PMF-powered ATP synthesis with different carbon sources and thus different growth conditions.

### The effect of F_1_F_o_ ATP synthase deletion on the growth rate and total yield depends on the carbon source

To test the strength of PMF and ATP coupling in different growth conditions, we compare fitness of the wild-type strain MG1655 (WT) with the full *atp* operon deletion mutant (referred to as Δ F_1_F_o_). ΔF_1_F_o_ strain lacks the PMF powered F_1_F_*o*_ ATP synthase/ATPase resulting in a complete uncoupling of PMF and ATP production. Fig. 2A shows growth curves of wild-type (grey) and Δ F_1_F_o_ (red) strains in M9 media with no casamino acids containing a single carbon source. Apart from glucose, we test disaccharides (lactose, maltose), pentose sugars (arabinose, xylose), non-fermentable carboxylic acids (acetate, succinate) and an alcohol (glycerol). Bacteria are grown in a 100-well honeycomb Bioscreen plate from 10^−6^ initial dilution of the overnight culture, see *Materials and Methods*. Figures 2B and C show the comparison of the growth rates and the maximum yield for the two strains. Fig. SI 3 shows the conversion of Bioscreen measured OD to the table top spectrophotometer with 1 cm optical path OD and to the cell number inferred from the CFU count.

**Fig. 2:**
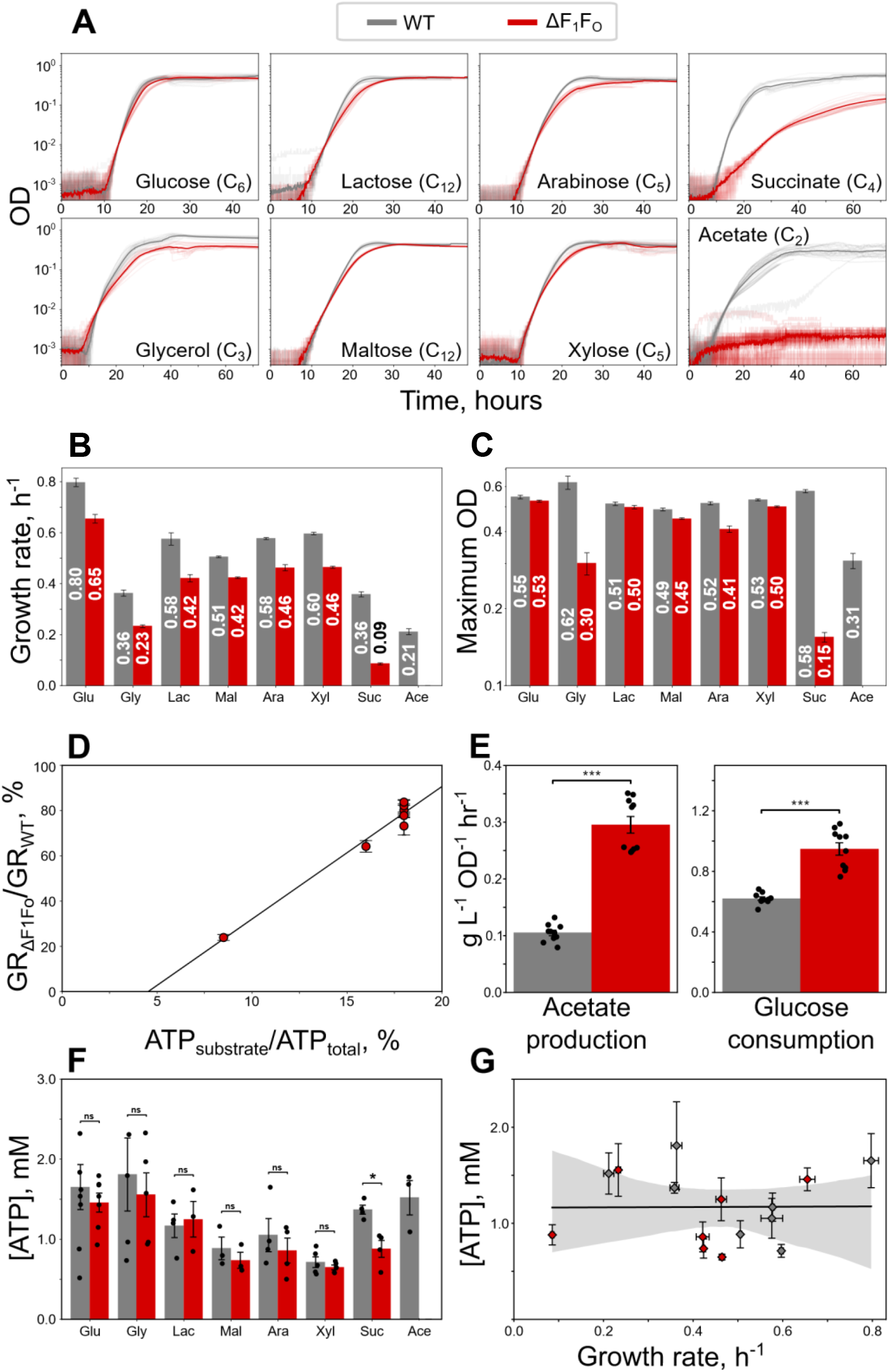
The effect of F_1_F_o_ deletion varies significantly depending on the available carbon source. Comparison of wild-type (grey) and ΔF_1_F_o_ (red) strains growing aerobically on different carbon sources. (**A**) Growth curves of wild-type and ΔF_1_F_o_ in minimal medium M9 supplemented with a single carbon source. Semi-transparent lines show individual curves, opaque lines are the means of the individual experiments (N varies from 13 to 33 repeats per condition). Individual curves are aligned at OD 0.01 for accurate averaging, except for acetate ΔF_1_F_o_ curves that never reach OD 0.01. Growth rates (**B**) and maximum OD (**C**) of wildtype and ΔF_1_F_o_ strains are extracted from the individual growth curves in (A) as described in *Materials and Methods*. The numbers on the bars show the mean of the individual growth rates/maximum OD, error bars are standard errors of the mean. (**D**) The ratios between the growth rates of ΔF_1_F_o_ and wild-type strains are correlated with the amount of ATP produced by substrate-level phosphorylation with respect to the total (maximum) ATP produced from a given carbon source. Grey line shows a linear fit. Pearson correlation coefficient of *r* = 0.932 indicates strong positive correlation between the variables. (**E**) The rates of acetate production and glucose consumption during aerobic growth on glucose is shown for wild-type and ΔF_1_F_o_ strains. Black points show individual measurements, the bars are means with standard errors. The difference between strains is statistically significant with a *t*-test *p*-value *p <* 0.001. (**F**) ATP concentration in the cytoplasm of cells grown on a single carbon source to OD 0.1 measured with luciferase assay, see *Materials and Methods* Black points show individual measurements, the bars are means with standard errors. (**G**) Growth rates and intracellular ATP concentrations are not correlated neither for wild-type nor for ΔF_1_F_o_ strains. The points and error bars are taken from the data on panels (B) and (C). The line is a linear regression with standard deviation shown as shaded region. Pearson correlation coefficient is equal to r = 0.008 indicating no correlation.

While using sugars (glucose, lactose, maltose, arabinose, or xylose) as a carbon source, ΔF_1_F_o_ performs surprisingly well reaching 72-82% of the wild-type growth rate and 78-98% of the wild-type total yield, indicating that loss of PMF-ATP coupling is not severely detrimental to *E. coli* in these conditions. Even on glycerol, commonly viewed as a poorer carbon source for significantly lower growth rates, ΔF_1_F_o_ strain reaches 64% of the wild-type growth rate and 48% yield. We note here, that loss of PMF-ATP coupling is not equivalent to purely fermentative anaerobic growth, and the F_1_F_o_ deletion strain aerobically reaches significantly higher yield than wild-type grown anaerobically [35]. The TCA cycle in ΔF_1_F_o_ strain in presence of oxygen can still run producing PMF as usual and ATP on the substrate level (1 ATP molecule per cycle turn), which allows the F_1_F_o_ deletion mutant to grow even on a non-fermentable carbon source succinate with 25% of the wild-type growth rate and yield (see Fig. SI 2 for the succinate metabolic pathway). Acetate, in contrast, yields zero net ATP on the substrate-level, making the ΔF_1_F_o_ strain growth impossible. Under these growth conditions, with no substrate-level ATP production, the PMF-ATP coupling becomes essential for *E. coli*.

We find that the reduction of the growth rate of the ΔF_1_F_o_ compared to the wildtype is correlated with the amount of ATP produced on a substrate level relatively to the theoretical maximum of ATP produced, Fig. 2D. The ATP yield calculations are given in Fig. SI 2 and SI Table 1. However, the apparent linear relation is not a direct translation of ATP_*substrate*_/ATP_*total*_ to the growth rate ratios, i.e., in conditions where substrate level ATP accounts for 12-23% of the theoretical total, the growth rate of the ΔF_1_F_o_ strain reaches 64-82% of wild-type growth rate. This could be explained by the increase in fermentation rate in the F_1_F_o_ deletion strain.

To estimate the increase in the rate of fermentation in the ΔF_1_F_o_ mutant compared to wild-type, which is needed to compensate for the loss of ATP synthase, we measure glucose consumption and acetate production in wild-type and ΔF_1_F_o_ cultures growing exponentially on glucose as the only carbon source, Fig. 2E. Acetate, produced by the phosphotransacetylase-acetate kinase (Pattack) pathway, is the main by-product of aerobic glucose fermentation in *E. coli* [12, 36, 37] and its accumulation in the medium can therefore serve as a measure of carbon flux through the fermentative pathway [13]. Acetate production rate for wild-type is 0.105 ± 0.005 g/L per OD unit per hour, and for ΔF_1_F_o_ 0.295 ± 0.015 g/L per OD unit per hour. Glucose consumption is 0.620 ± 0.012 g/L per OD unit per hour for wild-type and 0.948 ± 0.041 g/L per OD unit per hour for ΔF_1_F_o_. Taking acetate production rate as a proxy for the rate of fermentation, we calculate that to produce all of its ATP, ΔF_1_F_*o*_ mutant ferments 2.5-3 times more than the wild-type. Glucose consumption in the mutant is increased 1.4-1.6 fold compared to wild-type.

Next, we compare ATP levels in the cytoplasm of bacteria growing on different carbon sources with and without F_1_F_o_ present. We grow bacteria to OD = 0.1 maintained steadily in the bioreactor to measure ATP during exponential growth stage. Fig. 2F shows mean ATP concentration of the population of cells measured with luciferase assay, where each point shows one biological replicate. Although ATP concentration varies between carbon sources, the difference for each carbon source is not statistically significant between wild-type and ΔF_1_F_o_ strains (apart from succinate) with *p*-value of the two independent samples *t*-test *p >* 0.05. For all tested conditions cytoplasmic ATP concentration falls between 0.5 and 2.5 mM. In line with previous reports [38], and unlike other bacteria [39], in *E. coli* we did not find any correlation between the growth rate and the ATP concentration on the population level, Fig. 2G. The fact that deletion of F_1_F_o_ does not significantly affect the free ATP pool in the cell for most of the tested carbon sources implies that there is an optimal ATP concentration that *E. coli* aims to maintain irrespective of the means of ATP generation.

### F_1_F_o_ ATP synthase deletion becomes more detrimental if the carbon source is limited

In the previous section we explored the growth conditions where the carbon source is abundant and does not limit culture growth. Here, we compare the fitness of ΔF_1_F_o_ and wild-type strains growing on limited amounts of glucose.

We grow bacteria as before, in a 100-well Bioscreen plate, aerobically, at 37°C, in a minimal defined M9 medium supplemented with varying amounts of glucose, from 1 *µ*M to 30 mM, Fig. 3A. The growth rate and maximum OD comparison are shown in Fig. 3B and C. As expected, for both strains the growth rate stays constant down to 100 *µ*M glucose with ΔF_1_F_o_ maintaining 80-85% of the wildtype growth rate. However, with the decrease of the total produced biomass, the difference between the strains becomes more pronounced. Whereas at the highest tested glucose concentration of 30 mM ΔF_1_F_o_ reaches roughly similar if not slightly higher total biomass as the wild-type, at 1 mM it is only half of the wild-type biomass. While growth curves at lower concentrations become too noisy for a reliable analysis, we can conclude that for both strains growth becomes undetectable at the glucose concentrations between 1 and 10 *µ*M.

**Fig. 3:**
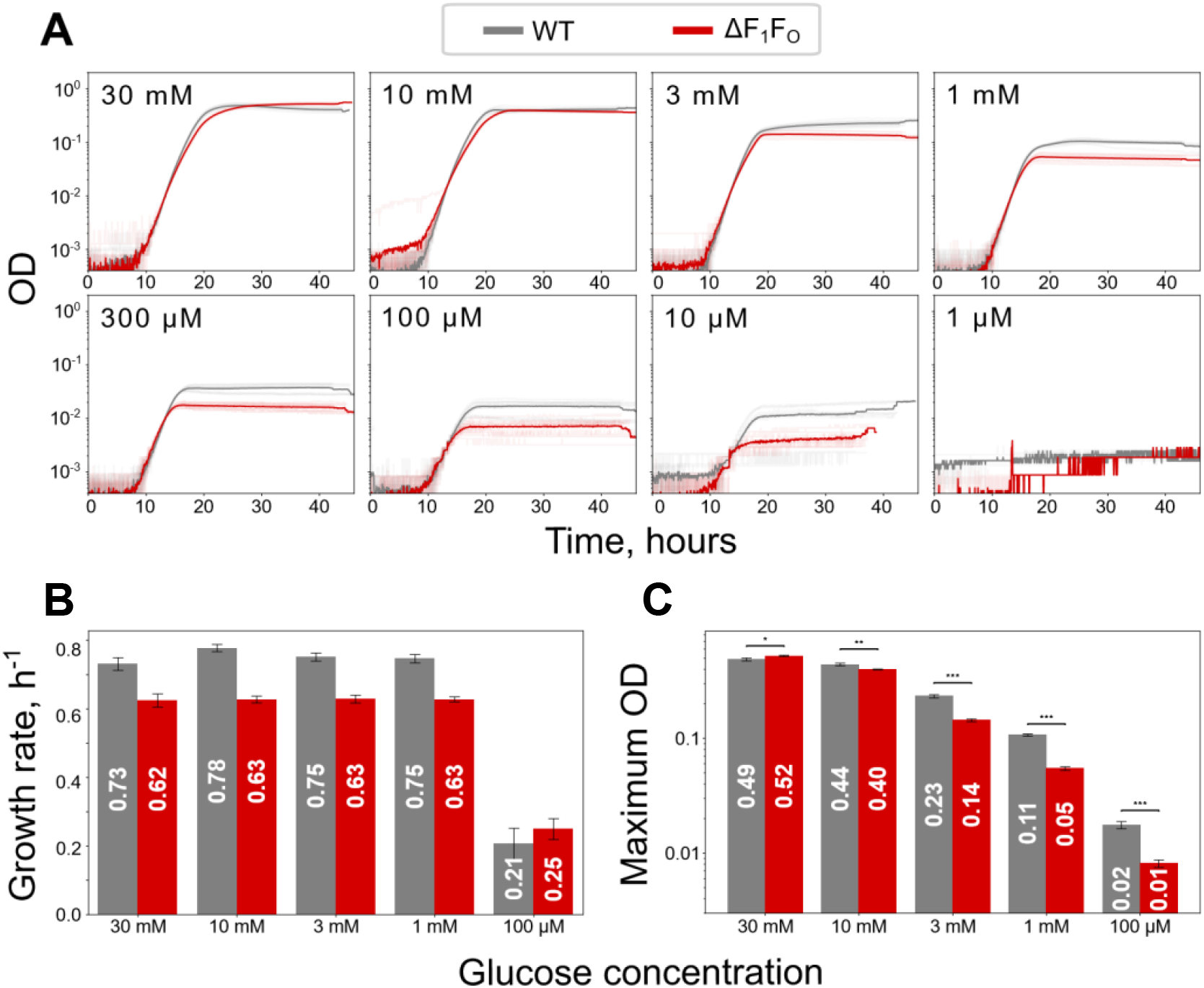
F_1_F_o_ deletion becomes important when the carbon source abundance is low. Comparison of wild-type (grey) and ΔF_1_F_o_ (red) strains growing aerobically on different concentrations of glucose as a single carbon source. (**A**) Growth curves of wild-type and ΔF_1_F_o_ in minimal medium M9 supplemented with a given concentration of glucose as a single carbon source in a Bioscreen plate reader. Semi-transparent lines show individual curves, opaque lines are the means of the individual experiments. Individual curves are aligned at OD 0.01 for accurate averaging where possible. Growth rates (**B**) and maximum OD (**C**) of wild-type and ΔF_1_F_o_ strains are extracted from the individual growth curves in (A) as described in *Materials and Methods*. The numbers on the bars show the mean of the individual growth rates/maximum OD, error bars are standard errors of the mean. The difference in maximum OD is statistically significant with (*) *t*-test *p*-value 0.01 *< p <* 0.05, (**) 0.001 *< p <* 0.01, and (***) *p <* 0.001.

These results demonstrate that, although F_1_F_o_ deletion affects *E. coli* growth on sugars only moderately, the effect becomes more significant when the carbon source abundance is limiting for growth.

### F_1_F_o_ is essential during anaerobic growth

So far, we have shown that F_1_F_o_ may be considered redundant during aerobic growth on sugars. We now ask how important the PMF-ATP coupling is for anaerobic growth.

Fig. 4 shows wild-type and ΔF_1_F_o_ growth curves on glucose or acetate in fully anaerobic conditions. We use acetate supplemented medium as an anaerobicity control, since *E. coli* is known to not be able to grow on acetate in the absence of oxygen or an alternative electron acceptor. Indeed, neither of the two strains grow on acetate confirming that the conditions are strictly anaerobic (see additional controls with glycerol and succinate in Fig. SI 4). On glucose, ΔF_1_F_o_ mutant fails to grow, while the wild-type grows as expected, confirming that F_1_F_o_ is essential during anaerobic growth [40].

**Fig. 4:**
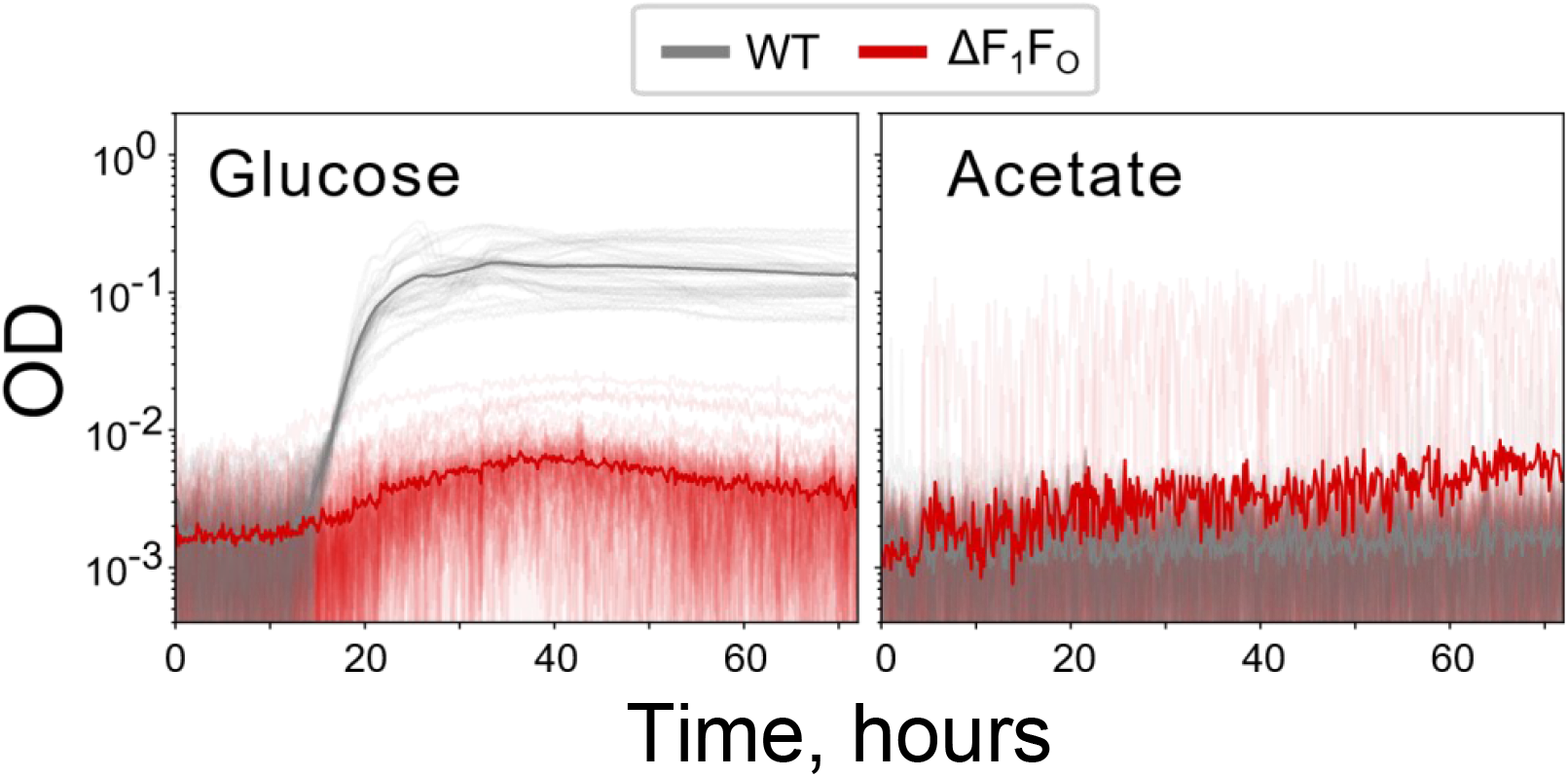
F_1_F_o_ is essential during anaerobic growth. Comparison of wild-type (grey) and ΔF_1_F_o_ (red) strains growing anaerobically in M9 supplemented with 0.3% glucose or acetate.

In purely fermentative mode *E. coli* can produce 4 ATP per molecule of glucose via substrate-level phosphorylation. However, in the absence of oxygen or other terminal electron acceptor, PMF is not being generated via ETC, since the last step of the process requires terminal electron acceptor reduction and re-oxidation of electron carriers NADH/FADH_2_. In the wild-type *E. coli* PMF can still be generated by ATP hydrolysis using ATPase function of the F_1_F_o_. But in the F_1_F_*o*_ deletion mutant this pathway does not exist. Although alternative pathways of PMF generation have been suggested in the literature [41, 42], they do not seem to be sufficient to produce enough PMF to support ΔF_1_F_*o*_ growth in fully anaerobic conditions where ETC is not functional. If the alternative electron acceptor (e.g., formate) is present in the medium, ΔF_1_F_o_ growth is partially restored Fig. SI 5, [42].

We therefore confirm that F_1_F_o_ is essential for anaerobic growth in *E. coli*. Moreover, unless we were to assume an alternative, yet unknown, role for F_1_F_o_, we can also safely conclude that PMF is essential for *E. coli* growth, irrespective of PMF’s role in ATP production.

## Discussion

The textbook view on cellular energy metabolism assumes that in the presence of oxygen ATP synthesis is mainly powered by PMF via the process of oxidative phosphorylation. Although generally not wrong, this view creates the perception that PMF is crucial for ATP production and, in turn, ATP synthesis is the main function of PMF. However, bacterial metabolism is complex and highly flexible, and is capable of diversification of ATP-generating pathways, which allows bacteria to optimise for speed [12], proteome and external resource allocation [13, 21, 22], membrane occupancy [43], and to adapt to various host environments [44]. On the other hand, PMF is essential for bacterial growth, even if it is not used for ATP synthesis [40]. Here, we estimated the strength of the PMF-ATP coupling in *E. coli* by assessing the fitness defect in bacteria when the coupling is fully removed by genetic deletion of the F_1_F_o_ ATP synthase/ATPase, and show that the degree of this coupling strongly depends on the growth conditions.

Under aerobic conditions, while growing on various sugars (glucose, lactose, maltose, arabinose, xylose) the fitness defect of the ΔF_1_F_o_ strain can be considered mild (less than 30% loss in growth rate, less than 25% yield reduction). It is consistent with previous reports [35, 45, 46], although we show a higher growth yield of the mutant strain than that in [45]. The difference has to do with the tested concentrations of carbon source. In [45] growth medium contains 1 mM glucose resulting in ΔF_1_F_o_ strain producing only half of the biomass of the wild-type, consistent with our results in Fig. 3. However, we show that at higher carbon source concentrations, ΔF_1_F_o_ can achieve similar, if not higher, total yield when compared to the wild-type.

Cells lacking PMF-ATP coupling achieve such high growth rates by re-routing their metabolic pathways towards PMF-independent ATP synthesis, increasing their fermentation rate by 2.5 to 3 fold compared to the wild-type. Although it causes less efficient use of the carbon source (∼ 50% higher glucose consumption), it does not affect the growth yield at high enough carbon source concentrations, where other factors, such as nitrogen abundance or metabolic waste accumulation, become limiting to the biomass increase. This showcases the adaptation potential of bacterial metabolism architecture, where a flux through one of the parallel pathways changes to compensate for the loss of function in the other.

The growth defect in the F_1_F_o_ deletion strain scales with the amount of ATP produced via substrate-level phosphorylation and becomes more significant with poor carbon sources such as glycerol and succinate. Acetate is an extreme example of a carbon source yielding zero net ATP at the substrate-level, making the PMF-powered ATP synthesis pathway essential for bacteria and fully inhibiting the growth of ΔF_1_F_o_ strain.

The other example of PMF-ATP coupling becoming essential is growth in anaerobic conditions. Here, the essentiality evidently comes from the inability to generate PMF via the electron transport chain in the absence of the terminal electron acceptor. In this case, proton efflux through F_1_F_o_ powered by ATP hydrolysis is a main source of PMF generation. This example emphasises the importance of the PMF outside of its role in ATP production. In the case of anaerobic growth on glucose, ATP is produced by fermentation, which means PMF is crucial for other aspects of the cell’s life, which cannot function in its absence. PMF is known to be involved in membrane transport [33, 47–49], motility [50–52], and pH maintenance [53]. Membrane voltage, one of the two components of PMF, was shown to be required for bacterial division in both Gram-positive (*Bacillus subtilis*) and Gram-negative (*E. coli*) bacteria [54]. The proposed mechanism behind it suggests an important role of the membrane voltage in the spatial organization of cytoskeletal and cell division proteins via modulation of membrane interaction with proteins. We dare to suggest that there might be other aspects of bacterial electrophysiology that still remain to be uncovered in order to fully explain the essential role of the PMF and membrane voltage in the bacterial life cycle.

We note that contradictory results on the permitting growth conditions of the oxidative phosphorylation mutant were reported in the earlier studies from the 1970’s [40, 55–58]. In [55] ATPase mutants were able to grow in anaerobic conditions, whereas in [40] they were not. In [58], out of multiple ATPase mutants tested, some were unable to grow on xylose, maltose, glycerol or succinate, while in [56] and [57] mutants growth on glycerol was shown. However, those studies relied on the selection, rather than genetic engineering, and could not account for other mutations in the strains, as they were done when DNA sequencing was just being invented. For example, it was later shown that growth of ATP mutants on succinate failed due to the transport efficiency rather than inability to produce ATP [59]. In this work we compare two strains with a precise genetic mutation that only differ by the ATP operon deletion.

An important consequence of the loose PMF-ATP coupling we report here is that the wild-type strain does not deplete its ATP pool upon losing PMF in glucose-rich medium, when cells are exposed to protonophores (CCCP, DNP) or electron transport chain/ATPase inhibitor (sodium azide). Although we did not measure the change in ATP/ADP ratio, which may better reflect the energetic state of the PMF-depleted bacteria, the gradual decrease of the ATP concentration and its high final level after the exposure to the uncouplers compared to that of cells in glucose-deficient medium suggest that ATP synthesis continues in the absence of PMF if the substrate for it is present. This result, consistent with a previously reported respiratory control in *E. coli* [60, 61], suggests that *E. coli* can switch to fully uncoupled ATP production within minutes if an appropriate carbon source is available. The dynamics do not change in a more complex medium, containing casamino acids, Fig. SI 6, although it causes more dramatic ATP depletion in a no glucose medium. We hypothesise that this is due to the remaining protein production, still possible in casamino acid-containing medium. As expected, CCCP causes no change in ATP concentration in the strain lacking F_1_F_o_ synthase in the glucose-rich medium (Fig. SI 7). However, surprisingly, in the medium with only trace amounts of glucose, ATP concentration in ΔF_1_F_o_ shows PMF-dependent drop, possibly, due to the voltage-powered membrane transport shutdown (Fig. SI 7).

Although ATP can be converted to PMF with F_1_F_o_ ATPase, when a protonophore is present, and therefore the electrochemical gradient across the membrane cannot be maintained, we observe that *E. coli* does not deplete its ATP pool on fruitless PMF production. It is not fully understood how F_1_F_o_ regulates the direction of its operation and prevents wasteful ATP hydrolysis. The known mechanisms of ATP hydrolysis inhibition include the IF_1_ protein [62–65] or ADP(Mg^2+^)-dependent inhibition [64–66]. There are also hypotheses that F_1_F_*o*_ exists in two different conformations: as an ATPase and as ATP synthase [64, 65, 67]. One possible explanation for the full ATP hydrolysis prevention in the presence of protonophores is that non-zero PMF is required to maintain F_1_F_o_ ATPase activity [63, 64].

F_1_F_o_ ATP synthase (and thus, PMF-ATP coupling) has been shown to be essential for some bacterial species: obligatory aerobes (*Mycobacterium tuberculosis, Acinetobacter baumannii, Mycobacterium smegmatis* [68, 69]) and obligatory or aerotolerant anaerobes (*Streptococcal* spp., *Clostridioides difficile, Lactococcus lactis* [68, 70]). For facultative anaerobes (*E. coli, Staphylococcus aureus, Listeria monocytogenes*) it is generally not considered essential [68], although for some, it is shown to become such in anaerobic conditions [40, 71, 72]. Here, we outline the conditions for its essentiality in *E. coli*, which could potentially apply to other facultative anaerobes.

The results of this work suggest a simplified view on bacterial energy metabolism of facultative anaerobes, Fig. 5. In this picture the accent is shifted from ATP as an ultimate energy output, to ATP and PMF, which are both required for bacterial growth. F_1_F_o_ here is presented as a bi-directional link between PMF and ATP, whose function is to ensure that both are produced in the cell, even if the critical components for their independent production are missing (i.e., ability to synthesise ATP on the substrate level or ability to oxidise NADH/FADH_2_ in the absence of a terminal electron acceptor). We believe that this simplified interpretation is instrumental for understanding of physiology and energy conversion in the bacterial cell not from the pure biochemical perspective but in the light of molecular and systems biology.

**Fig. 5:**
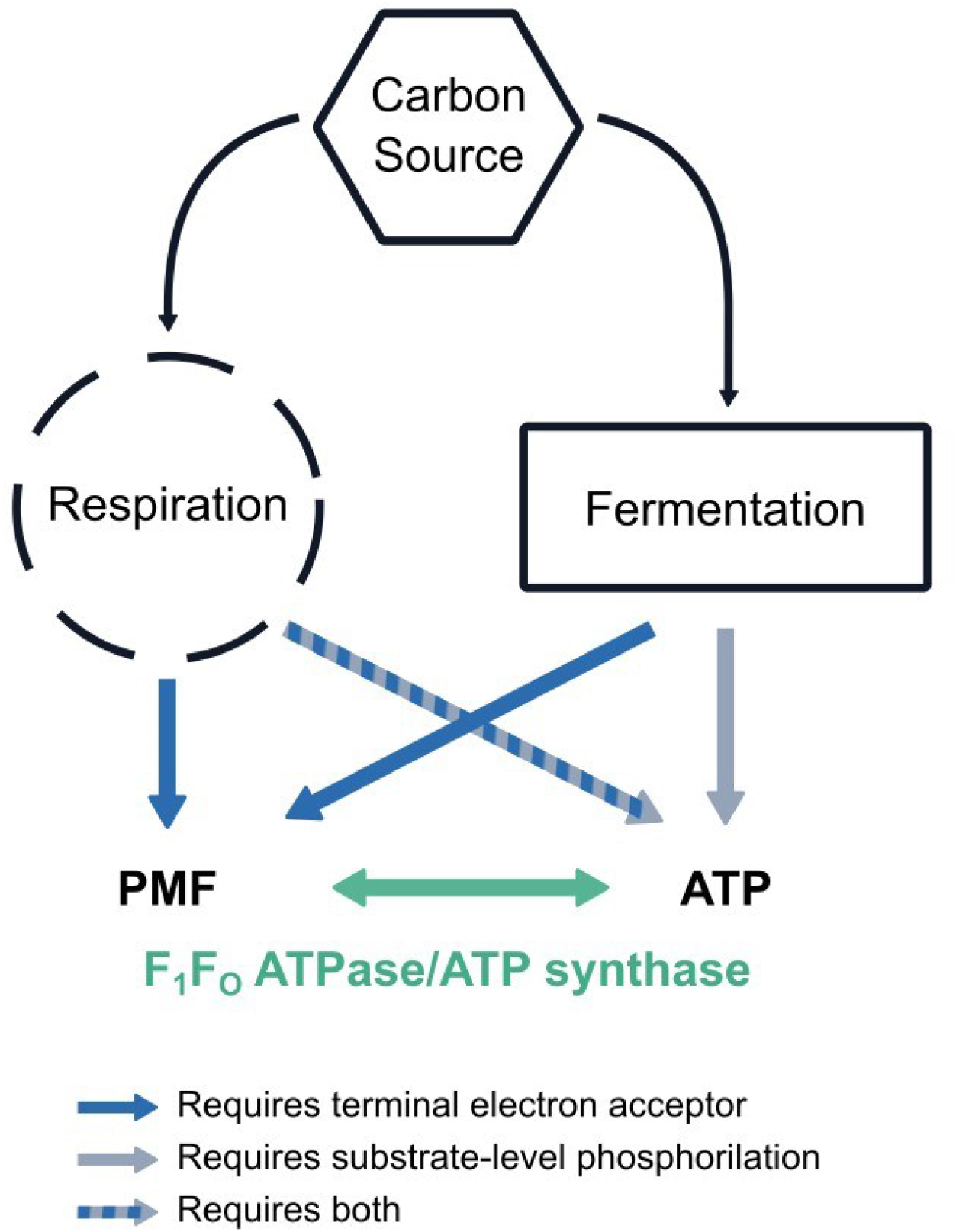
*E. coli* energy metabolism. PMF and ATP are produced by both fermentative and respiratory pathways if the conditions for their independent generation are met. The F_1_F_o_ ATPase/ATP synthase plays a role of bi-directional link providing an auxiliary pathway for the PMF and ATP generation if conditions are satisfied only partially.

## Materials and Methods

### Strains and media

K-12 MG1655 was used as a wild-type. ΔF_1_F_o_ was obtained by removing full *atp* operon, *atpC*-*atpI*, from wild-type strain by a plasmid mediated gene replacement method [73]. For upstream homology arm we used primers preC for 3’ - CCAGGTGAACGTGCGCTG and preC rev 3’ - CACCGGCTTGAAAAGCACAAAAG, for downstream homology postI for 3’ - CACGTTTTTCACTCCTGCTCCC and postI rev 3’ - ACTCTTTTGCATCAACAAGATAACGTG. The entire region between preC rev and postI for was deleted from the MG1655 chromosome and the deletion was verified with the full-genome sequencing.

For most of the experiments we used defined M9 minimal medium (5x M9 minimal Salts pH 7.2, M6030, Sigma-Aldrich; 2 mM Mg_2_SO_4_; 0.1 mM CaCl_2_) supplemented with 0.3% of a given carbon source, unless stated otherwise. pH was not adjusted extra and ranged from 7.2 to 7.5 for most media, apart from succinate with pH 6.1. For the experiments in Fig. 1, M9 medium was also supplemented with 0.2% casamino acids from the autoclaved 2% stock solution (Casein hydrolysate, Roth AE41.1). For overnight cultures we used Lysogeny Broth (LB: 10 g tryptone, 5 g yeast extract, 10 g NaCl per 1 L, pH 7.0).

### Growth curves

#### Pre-culture

For the plate reader experiments, single colonies of wild-type and ΔF_1_F_o_ strains from the fresh plates were inoculated into 2 ml LB in a culture tube and grown overnight at 37°C with shaking at 220 rpm. The resulting OD was OD_*WT*_ ≈ 5.0 and 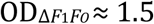.

#### Aerobic growth

Growth curves in Fig. 2 and 3 were measured with Bioscreen C reader (Oy Growth Curves Ab Ltd., Finland) in a 100-wells honeycomb plates in 300 *µ*l culture volume from 10^−6^ for ΔF_1_F_o_ and 2·10^−6^ for wild-type strain dilution of the pre-culture at 37°C with shaking. Readings were taken every 4 minutes with fast shaking at maximum amplitude stopping 5 seconds prior to 600 nm absorption measurement.

OD calibration between the Bioscreen, table-top spectrophotometer (WPA CO 8000 Cell Density Meter, Biochrom, UK), and the cell number is given in Fig. SI 3.

#### Anaerobic growth

For anaerobic growth (Fig.4, SI 4, and SI 5) we used anaerobic chamber (gas mixture: 85% N_2_, 10% CO_2_, 5% H_2_). The growth media bottles with partly unscrewed caps were kept in the chamber for at least 12 hours before the experiment. Aerobic pre-cultures in LB were taken inside the chamber, and diluted 10^−6^ or 2·10^−6^ into 200 *µ*l of growth media. The measurements (600 nm absorption) were taken with MultiscanTM GO plate reader (Thermo Fischer Scientific, USA) installed inside the anaerobic chamber, every 10 minutes with continuous shaking at 37°C in transparent flat bottom 96-well Nunc™ plates (Thermo Fischer Scientific, USA).

#### Growth curves analysis

Growth curves were analysed with custom written python scripts, using *pandas, numpy* and *scipy* packages. For each curve, the mean OD value for the first 100 minutes (25 measurements) was taken as a blank and subtracted from the remaining growth curve. A fixed window of OD values between 0.005 and 0.05 (in Bioscreen OD scale) was chosen for exponential function (*OD* = *OD*_0_ ·*e*^*µt*^, where *OD*_0_ is initial OD, *µ* is a growth rate, and *t* is time in hours) fitting for all conditions. An example of curve fitting is given in Fig. SI 8. The growth rates and maximum OD values were calculated for each individual curve and the mean and standard error was then calculated for each condition.

### ATP measurements

ATP was measured on the population level using BacTiter-Glo™ luciferase assay (G8232, Promega, USA) at room temperature according to the manufacturer instruction. For luminescence measurements we used black Nunc™ 96-Well OpticalBottom Microplates (Thermo Fischer Scientific, USA). 100 *µ*l of the bacterial culture was mixed with 100 *µ*l of BacTiter-Glo™ reagent, shaken in the plate reader for 30 sec at 807 cpm (1 mm amplitude), and incubated for another 4.5 minutes. Endpoint luminescence was then measured by Synergy™ H1 plate reader (BioTek Instruments, Inc., USA) with 170 gain. The ATP calibration was performed as per manufacturer instruction: 100 mM ATP stock solution (Invitrogen™) was diluted in a 10-fold series and luminescence was measured in the interval between 0.1 nM to 10 *µ*M. Later, for each experiment a 2-point (100 nm and 1 *µ*M) or a 3-point (10 nm, 100 nm, and 1 *µ*M) calibration was performed as a control.

#### Dynamic ATP measurements

For ATP dynamics measurements (Fig. 1, Fig. SI 6, and Fig. SI 7) bacterial cultures were grown in M9 medium supplemented with 0.3% glucose (and 0.2% casamino acids for Fig. SI 6 and Fig. SI 7) in an OGI BioReactor (Ogibiotech, UK) in a turbidostat mode maintaining constant OD 0.4 at 37°C. 5 ml of the steady state exponential culture was then transferred to a culture tube, cooled down to the room temperature (according to the BacTiter-Glo™ instructions) and a zero point ATP concentration (prior to the treatment) was measured. 100 *µ*M of CCCP (or 5 mM of sodium azide, or 2.5 mM DNP) was then added to the culture and ATP was measured every 5-10 minutes for the total 1 hour. CFU counts were taken before and after the treatment. For CFU counts the culture was diluted to 10^−5^ and 10^−6^ and 100 *µ*l of each dilution was plated onto LB agar plates with no selection. The colonies were counted after overnight growth at 37°C.

#### Single point ATP measurements

For a single point ATP measurements the cultures were grown in a corresponding media (M9+0.3% of a given carbon source) in OGI BioReactor (Ogibiotech, UK) in a turbidostat mode to OD 0.1 (to measure ATP in an early exponential culture at the highest growth rate). The luciferase measurements were done as described above with 3 technical replicates for each biological replicate and CFU counts performed with 10^−4^, 10^−5^, and 10^−6^ dilution of the bacterial culture.

#### Cell sizes measurements

To convert bulk ATP measurements to single cell ATP concentrations we measured the cell sizes in all the tested conditions with phase contrast microscopy. Cells culture was taken from the turbidostat (growing at constant OD 0.4 or 0.1), concentrated 10x, and added to the glass slide. Phase contrast images were taken with Nikon Eclipse Ts2R microscope, with 100x Nikon Ph3 DL oil objective. The length and the width of 30-100 cells per strain per condition were measured in Nikon NIS-Elements D software. The measured length (*L*) and width (*W*) were converted to cell volumes, assuming *E. coli* as a spherocylinder (or capsule), as follows: 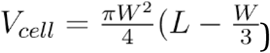. Sizes distributions are shown in Fig. SI 1.

### Acetate production/glucose consumption measurements

For acetate production and glucose consumption measurements wild-type and ΔF_1_F_o_ strains were grown from single colonies overnight in 2 ml M9 supplemented with 0.3% glucose in culture tubes. The overnight cultures were then diluted 1:1000 into 50 ml of fresh M9+0.3% glucose and grown in 250 ml conical flasks at 37°C with shaking at 220 rpm. OD was measured every 1 hour with table-top spectrophotometer (WPA CO 8000 Cell Density Meter, Biochrom, UK). Media samples were taken at OD range 0.5-1 (table-top spectrophotometer scale, see Fig. SI 9 for actual growth curves) by sterilising 1.5 ml of culture with 0.22 *µ*m syringe filter. Samples were frozen and kept at -20°C until measurements. To measure initial concentrations of acetate (*c*_*a*0_ = 0 g/L) and glucose (*c*_*g*0_ = 2.85 ± 0.04 g/L) in the medium, samples of pure M9+0.3% glucose was taken.

Glucose and acetate concentrations were quantified using a Vanquish HPLC system (Thermo Fisher Scientific, USA) equipped with an Aminex® HPX-87H column (300 × 7.8 mm; Bio-Rad Laboratories, USA) and a refractive index detector (RefractoMax 520; ERC Inc., Japan). The mobile phase consisted of 4 mM H_2_SO_4_ prepared in Milli-Q water, operated at a flow rate of 0.5 mL/min. Samples were once more filtered through 0.20 *µ*m Mini UniPrep Syringeless PES filters (Whatman). Quantification was performed using external calibration curves generated from glucose standards (1, 5, 10, 25, and 50 g/L) and acetate standards (0.01, 0.05, 0.10, 0.25, 0.50, and 1.00 g/L). The chromatographic method was run at 30°C for 30 min with an injection volume of 1 *µ*L per standard or sample.

Such obtained concentrations of glucose (*c*_*g*_) or acetate (*c*_*a*_) were converted to production/consumption rates (*R*) as follows: 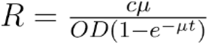, where *µ* is a growth rate, *OD* is a culture OD at which the sample was taken, *t* - time from inoculation in hours, and *c* = *c*_*a*_ for acetate production, or *c* = *c*_*g*0_−*c*_*g*_ for glucose consumption.

## Supplementary Material

### Supplementary Tables

**Table 1:**
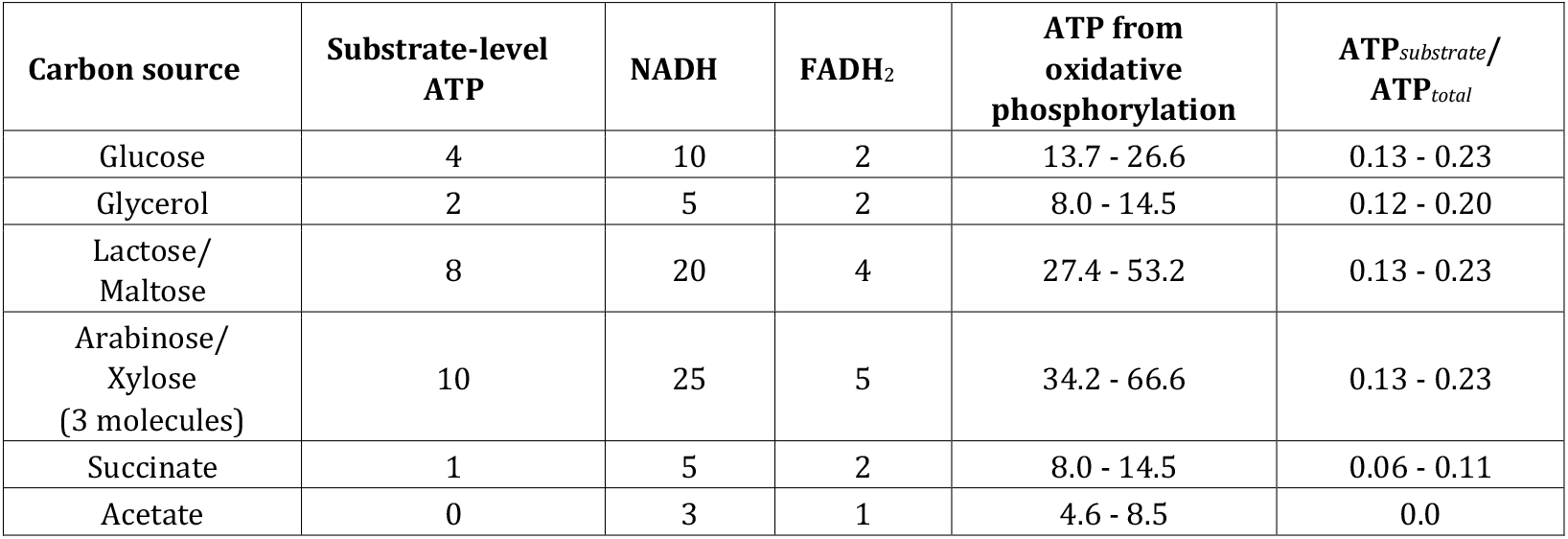
Amount of ATP produced via substrate-level or oxidative phosphorylation on different carbon sources. To calculate ATP yield from NADH and FADH_2_, we use the following estimates: 1 NADH molecule can transport 4-8 H^+^ across the membrane, 1 FADH_2_ - 4 H^+^, while F_1_F_o_ ATP synthase uses 3.33.5 H^+^ for production of 1 molecule of ATP [74]. Thus, we take NADH:ATP ratio as 1:1.14-1:2.42 and FADH_2_:ATP as 1:1.14-1:1.21. For arabinose/xylose pathways the output is given for 3 input molecules. For the detailed pathways see Fig SI 2. GTP is considered ATP equivalent.

| Carbon source | Substrate-level ATP | NADH | FADH<sub>2</sub> | ATP from oxidative phosphorylation | ATP<sub>substrate</sub>/ATP<sub>total</sub> |
| --- | --- | --- | --- | --- | --- |
| Glucose | 4 | 10 | 2 | 13.7 - 26.6 | 0.13 - 0.23 |
| Glycerol | 2 | 5 | 2 | 8.0 - 14.5 | 0.12 - 0.20 |
| Lactose/<br>Maltose | 8 | 20 | 4 | 27.4 - 53.2 | 0.13 - 0.23 |
| Arabinose/<br>Xylose<br>(3 molecules) | 10 | 25 | 5 | 34.2 - 66.6 | 0.13 - 0.23 |
| Succinate | 1 | 5 | 2 | 8.0 - 14.5 | 0.06 - 0.11 |
| Acetate | 0 | 3 | 1 | 4.6 - 8.5 | 0.0 |

### Supplementary Figures

**Figure SI 1.**
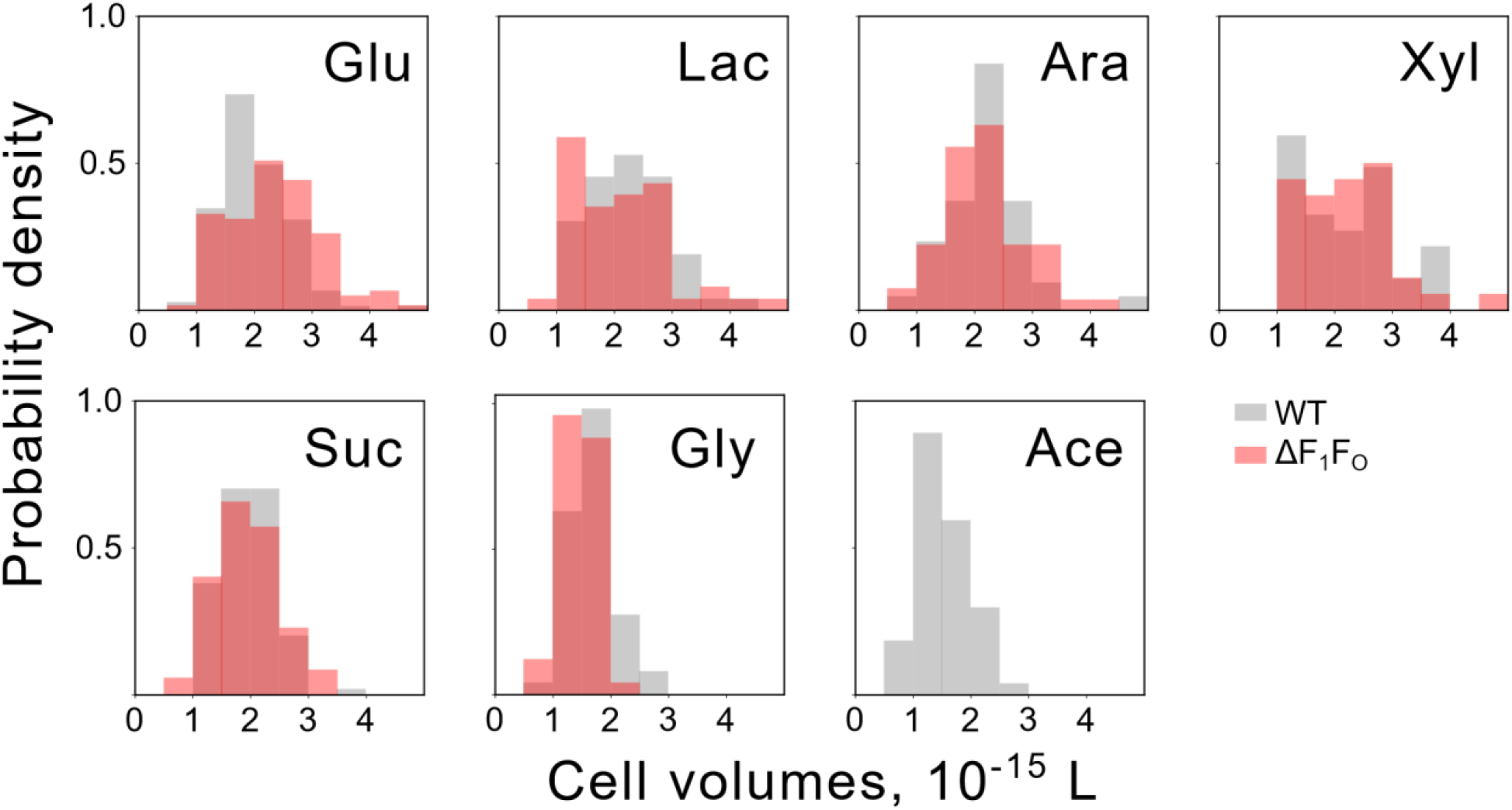
Distributions of wild-type and ΔF_1_F_o_ cell sizes on different carbon sources. Cell cultures were grown in turbidostat at OD 0.1. For ATP concentration calculations, cell sizes are measured as described in *Materials and Methods*. The sizes of 36 - 150 cells per strain per conditions were analysed. For sugar carbon sources, mean volume for both wild-type and ΔF_1_F_o_ cells was taken as V_*cell*_=2.2·10^−15^ L, for succinate V_*cell*_=2.0·10^−15^ L, for acetate and glycerol V_*cell*_=1.5·10^−15^ L.

**Figure SI 2.**
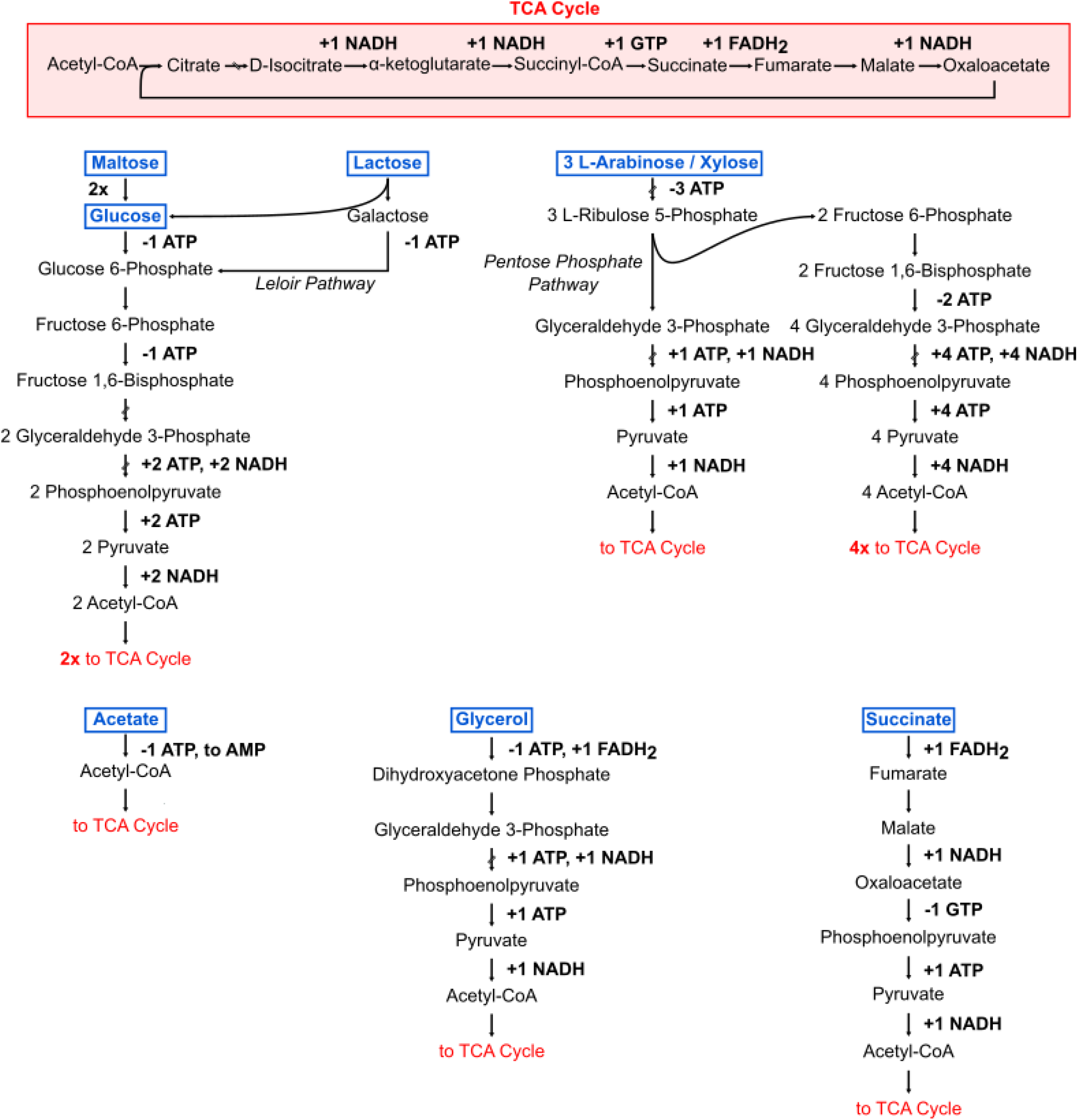
Simplified metabolic pathways for various carbon sources in aerobic conditions. Red box shows production of GTP, NADH, and FADH_2_ within tricarboxylic acid (TCA) cycle. Below, carbon sources (blue) are shown to be metabolized, producing ATP, GTP, reducing agents (NADH, FADH_2_), and Acetyl-CoA. Acetyl-CoA is then fed into the TCA cycle. For maltose “2x” mark next to an arrow means the duplication of all the downstream reactions and products. Both lactose and maltose produce double the yield of glucose, since maltose gets broken into two glucose molecules, and lactose – to glucose and galactose, which later enters the same glucose pathway via Leloir pathway. Intermediates that do not function as nodes between pathways are skipped with an interrupted arrow for clarity. For arabinose/xylose pathway input requires three molecules of the sugar. Production and consumption of ATP is a conversion with ADP and P_*i*_, unless otherwise stated.

**Figure SI 3.**
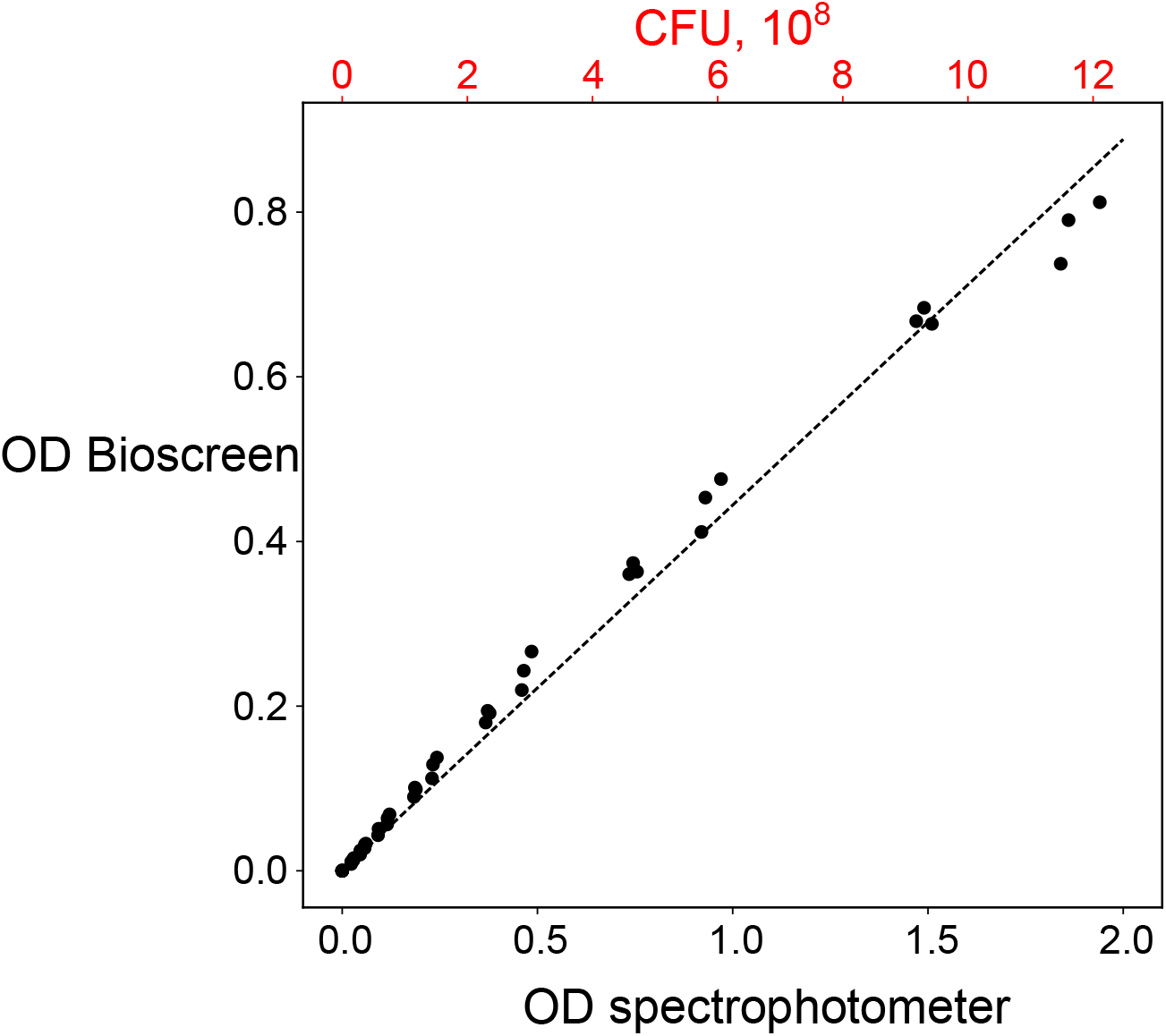
Bioscreen OD conversion to table-top spectrophotometer OD and to CFU. 3 overnight cultures of each wild-type and ΔF_1_F_o_ strains were serially diluted and OD was measured in a Bioscreen plate (300 *µ*l sample volume) and in a standard cuvette of the table top spectrophotometer WPA CO 8000 Cell Density Meter with 1 ml samples. Initial cultures diluted to 10^−5^, 10^−6^, and 10^−7^ were plated for cell count and OD conversion to CFU. The mean for 6 cultures of 6.24·10^8^ CFU/ml/OD was calculated. The dotted line shows linear fit *OD*_*Bio*_ = *aOD*_*Sph*_, where *a* = 0.44.

**Figure SI 4.**
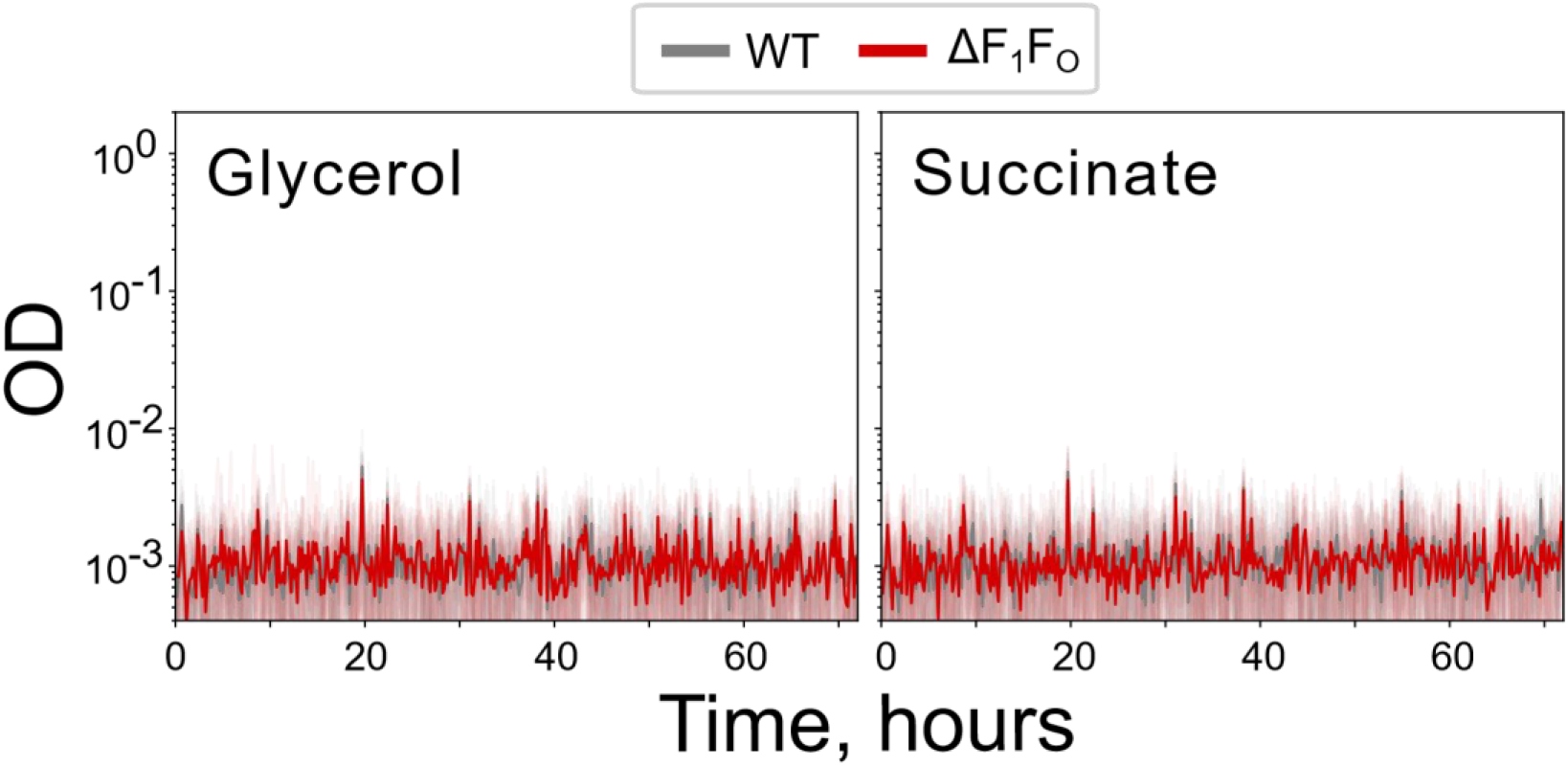
Neither wild-type nor ΔF_1_F_o_ strains grow anaerobically on glycerol or succinate without an alternative electron acceptor. Growth protocol in the anaerobic chamber is the same as in the Main text Figure 4, see *Materials and Methods*.

**Figure SI 5.**
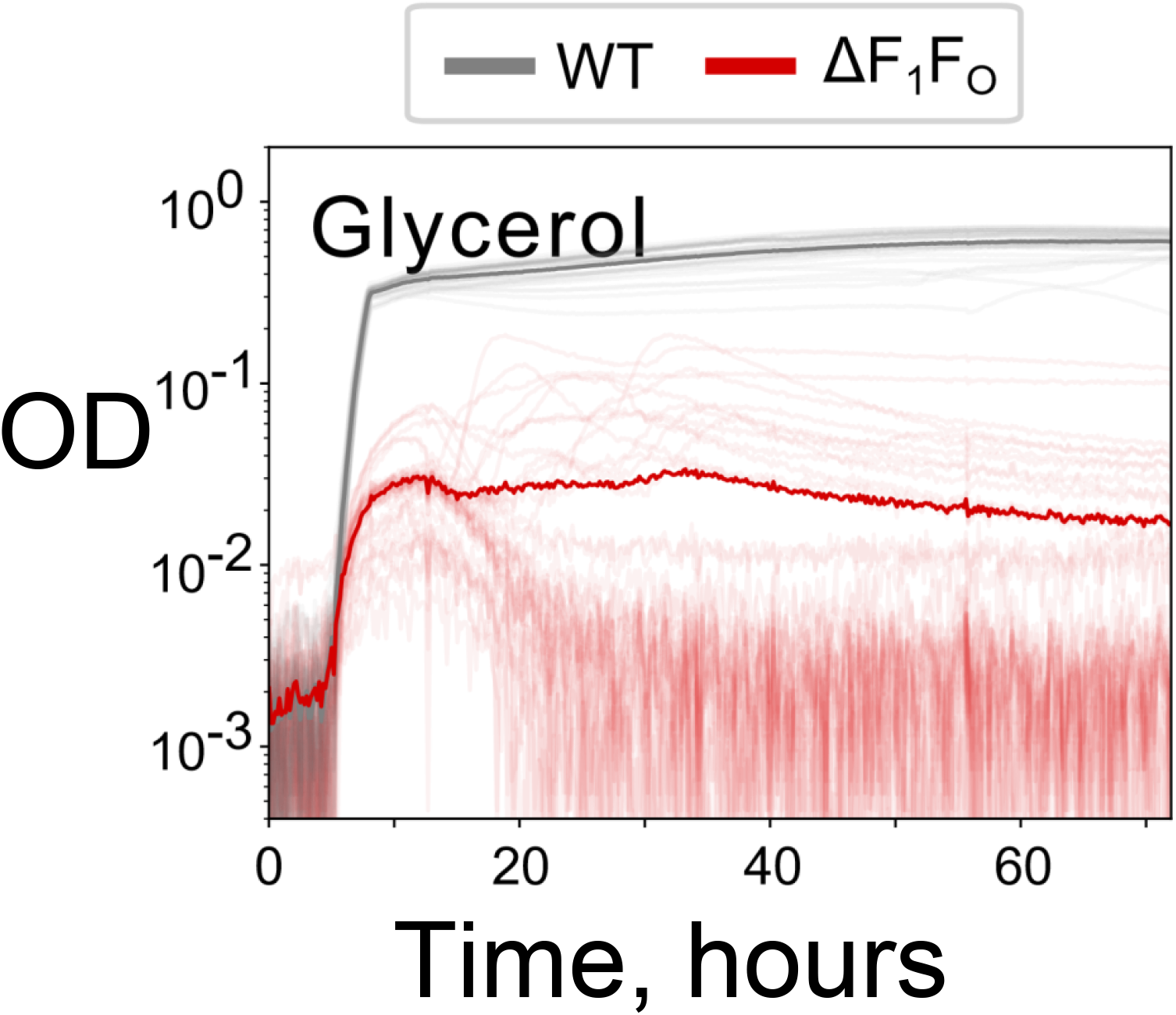
ΔF_1_F_o_ strains growth is possible in anaerobic conditions if an alternative electron acceptor (e.g., formate) is present. wild-type and ΔF_1_F_o_ strains are grown in an anaerobic chamber in mixed acid medium reproduced from [42] (20 g/L peptone, 15 g/L K_2_HPO_4_, 1.1 g/L KH_2_PO_4_, 5 g/L NaCl, 11.1 mM glucose, 137 mM glycerol, and 10 mM sodium formate). All of the ΔF_1_F_o_ cultures had an initial spike of growth reaching OD∼0.05, after which one third of the cultures (12 out of 36) continued to grow, while the rest collapsed. Only a few cultures reached OD 0.1. wild-type strain growth was consistent for all biological and technical replicates.

**Figure SI 6.**
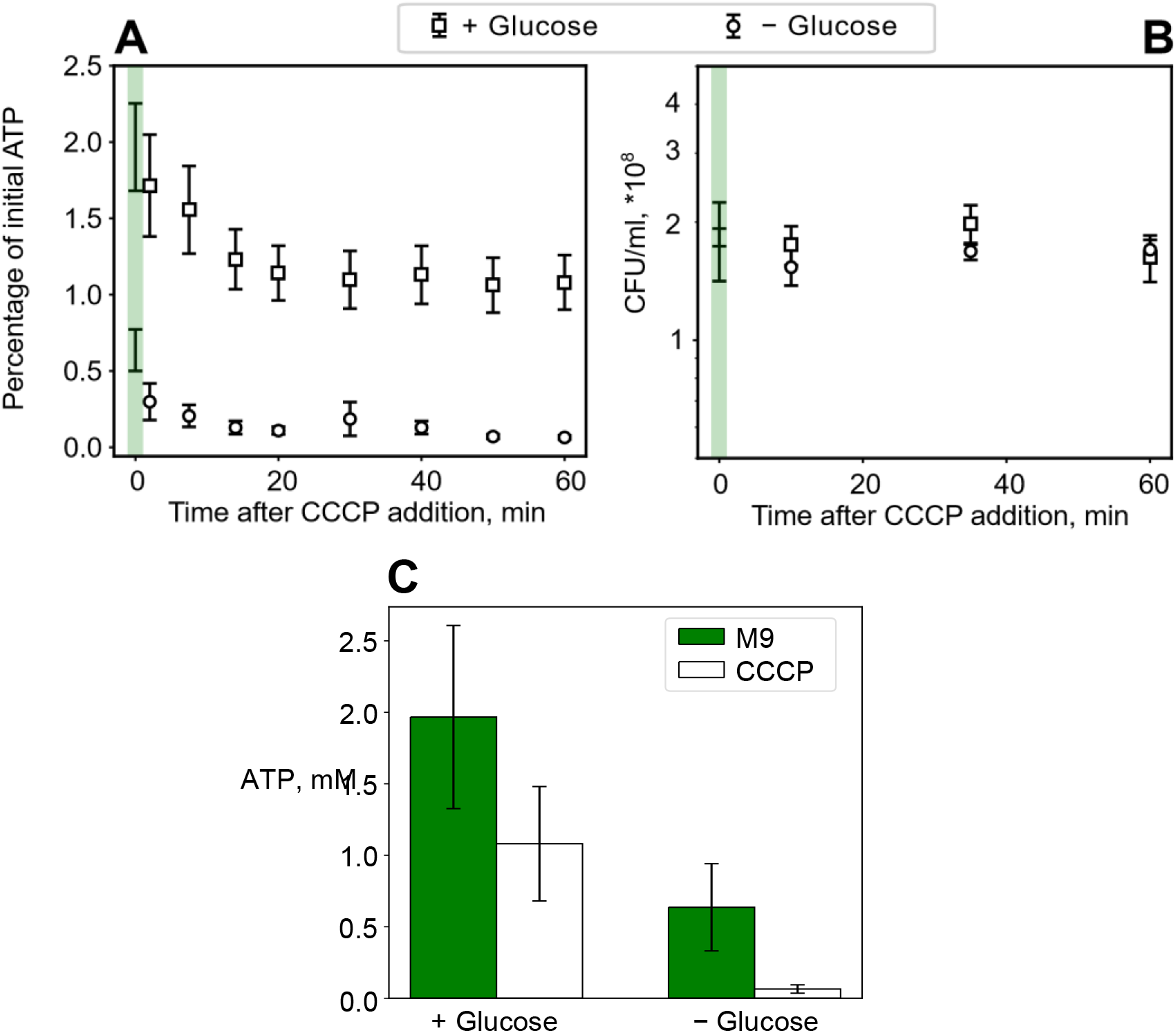
CCCP does not fully dissipate ATP in wild-type MG1655 if glucose is present in the complex M9 medium containing casamino acids. (**A**) Time series of ATP concentration in *E. coli* MG1655 cytoplasm upon addition of 100 *µ*M CCCP in M9 medium containing 0.2% casamino acids and either 0.3% (squares) or no glucose (circles). As in the main text Fig. 1, CCCP is added at time t=0 and ATP concentration is measured every 5-10 minutes for a total of 1 hour with luciferase assay. The first point at t=0 shows ATP concentration prior to treatment (highlighted by the green shading). Error bars show standard deviation of the mean for N=6 independent experiments. (**B**) CFU counts before (t=0) and during (t=10, 30, or 60 min) the treatment. Within 60 minutes of the experiment CFU count stays constant irrespective of glucose. (**C**) Absolute ATP concentration in cells’ cytoplasm before (t=0, green bars) or after (t=60 min, white bars) CCCP addition, with and without glucose. In presence of glucose ATP concentration changes from 1.97 ± 0.29 mM to 1.08 ± 0.18 mM, in the medium devoid of glucose, initial ATP concentration of 0.64 ± 0.14 mM drops to 0.06±0.01 mM.

**Figure SI 7.**
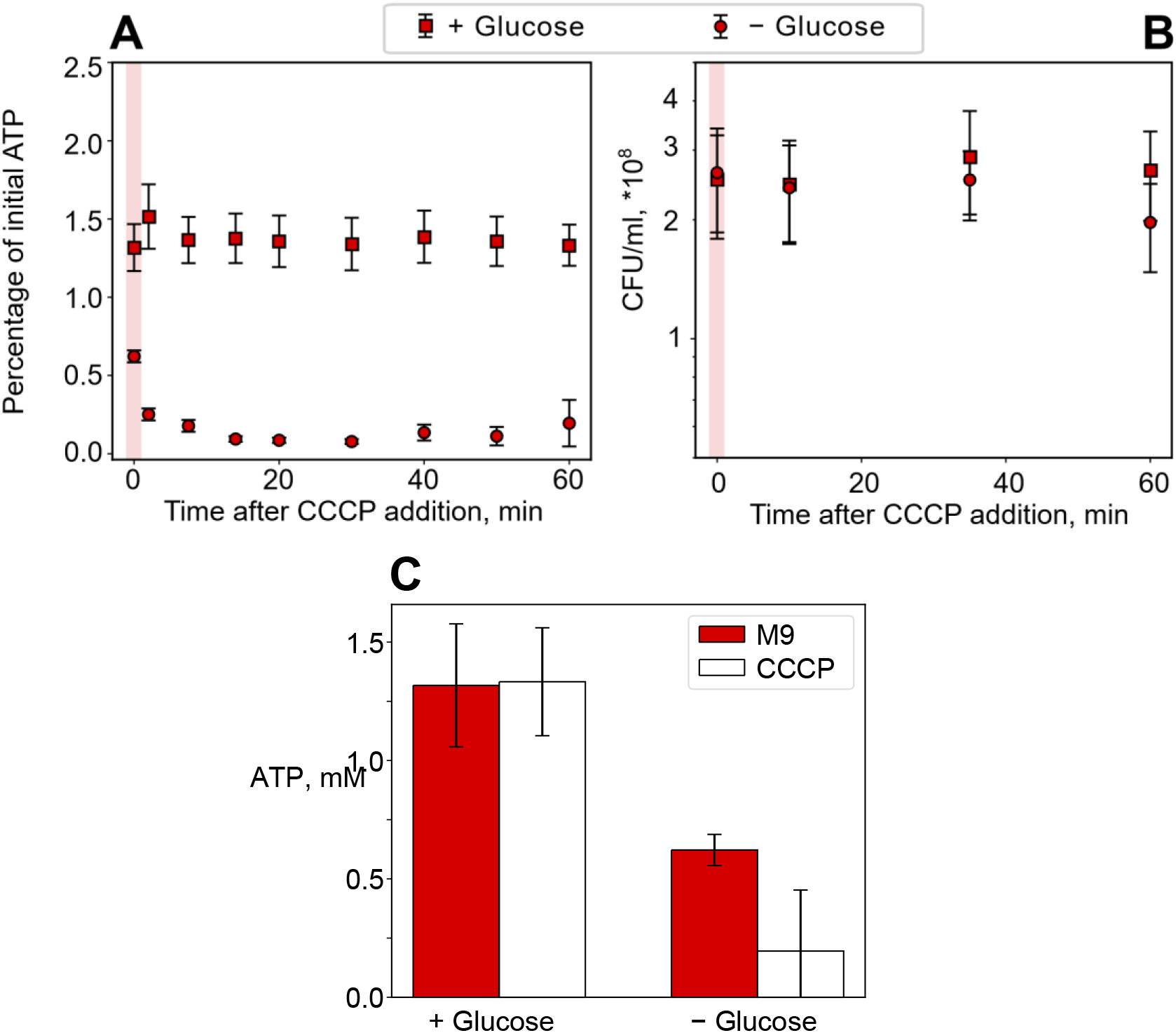
CCCP does not change ATP concentration in the ΔF_1_F_o_ cells if glucose is present in the medium. ΔF_1_F_o_ cells were treated the same way as wild-type cells in Fig. SI 6. Briefly, cell culture in M9+0.2% casamino acids+0.3% glucose grown to OD 0.4, was washed into either the same medium containing 0.3% glucose (squares), or to the same medium but with no glucose added (circles), and treated with 100 *µ*M CCCP. (**A**) ATP concentration was measured with the luciferase assay every 5-10 minutes for 1 hour. Red shading shows ATP level measured prior to CCCP addition (t=0). As expected, ΔF_1_F_o_ strain did not change ATP concentration upon CCCP addition in the glucose-rich medium, since in ΔF_1_F_o_ strain ATP is produced completely independently of PMF. Surprisingly, in glucose-deficient medium the ATP level dropped significantly even in the absence of F_1_F_o_. We hypothesise that the drop is related to the glucose transport rather than glucose metabolism in a poor medium. While in the glucose-rich environment glucose transport does not require PMF, in a medium with trace amounts of glucose, voltage-powered membrane transport may play a significant role in cells’ metabolism. (**B**) Similarly to wild-type, exposure to CCCP does not cause cell death in ΔF_1_F_o_ strain within 1 hour of experiment. (**C**) ATP concentration before (t=0, red) and after (t=60 min, white) CCCP addition in the glucose-rich and glucose-deficient media. Error bars show standard deviation for N=4 experiments.

**Figure SI 8.**
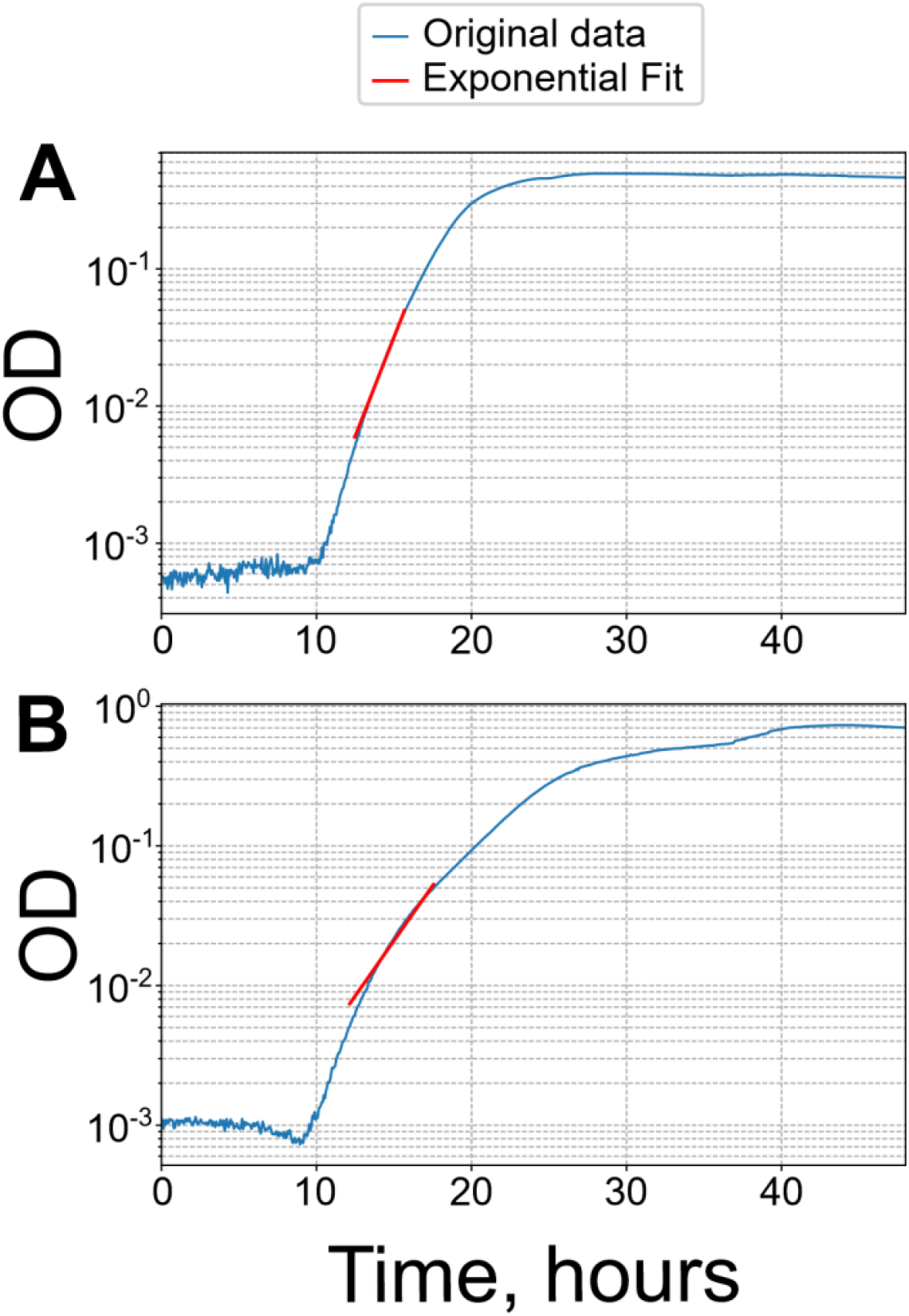
Example of fit procedure for growth rate calculation. A fixed OD window of 0.005 to 0.05 was chosen for exponential fitting as a reasonable loglinearity range for both fast-growing (glucose, **A**) and slow-growing (glycerol, **B**) cultures.

**Figure SI 9.**
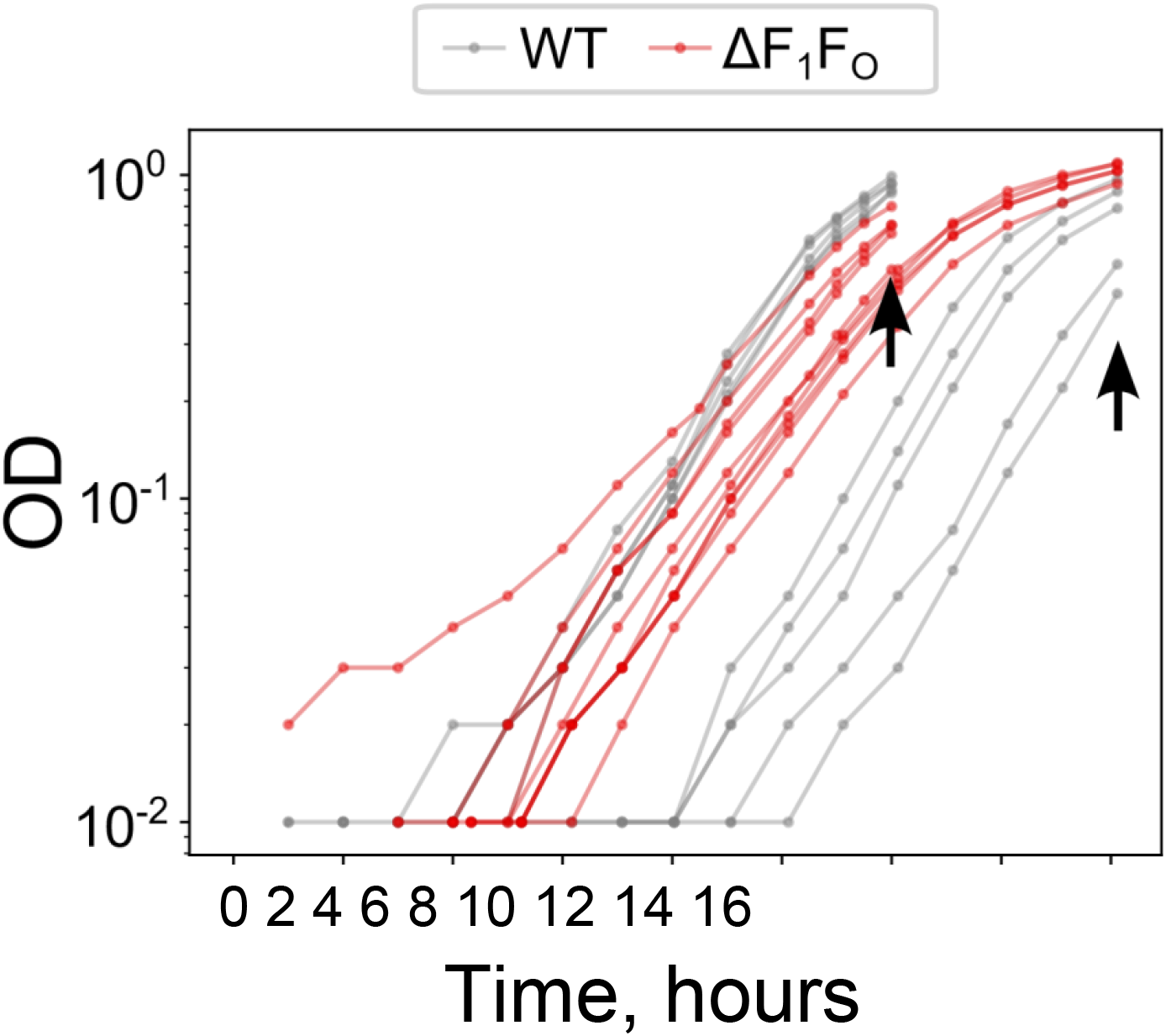
Growth curves for samples analysed for acetate consumption/glucose production rates. The sample collection for acetate production and glucose consumption rates was done on two different days. Each day, 5 biological repeats of wild-type and ΔF_1_F_o_ strains were grown in 50 ml of M9 supplemented with 0.3% glucose in 500 ml conical flasks with shaking at 37°C for 12-16 hours to OD 0.5-1. Measurements were taken with table-top spectrophotometer WPA CO 8000 Cell Density Meter every hour. Arrows indicate points of supernatant samples collection.

